# Polyanions provide selective control of APC/C interactions with the activator subunit

**DOI:** 10.1101/715615

**Authors:** Arda Mizrak, David O. Morgan

## Abstract

Transient interactions between the Anaphase-Promoting Complex/Cyclosome (APC/C) and its activator subunit Cdc20 or Cdh1 generate oscillations in ubiquitination activity necessary to maintain the order of cell cycle events. Activator binds the APC/C with high affinity and exhibits negligible dissociation kinetics *in vitro*, and it is not clear how the rapid turnover of APC/C-activator complexes is achieved *in vivo*. Here, we describe a mechanism that controls APC/C-activator interactions based on the availability of substrates. We find that APC/C-activator dissociation is stimulated by abundant cellular polyanions such as nucleic acids and polyphosphate. Polyanions also interfere with the ubiquitination of low-affinity substrates. However, engagement with high-affinity substrates blocks the inhibitory effects of polyanions on activator binding and APC/C activity. This mechanism amplifies the effects of substrate affinity on APC/C function, stimulating processive ubiquitination of high-affinity substrates and suppressing ubiquitination of low-affinity substrates.

## INTRODUCTION

Regulatory systems in biology often depend on the transient formation of specific protein complexes. Dynamic interactions between protein partners must be achieved in the crowded and complex environment of the cell, where countless nonspecific collisions with other macromolecules can influence the rate of complex formation and disassembly. Numerous mechanisms are employed to enhance the specificity of protein interactions under these conditions, but our understanding of this problem is limited in part because studies of these systems often depend on analysis with purified components.

Dynamic and specific protein-protein interactions lie at the heart of the regulatory system that governs progression through the cell division cycle. One of the key components of this system is a ubiquitin ligase, the Anaphase-Promoting Complex/Cyclosome (APC/C), which triggers chromosome segregation and the completion of mitosis^1,2^. The APC/C is activated at specific cell-cycle stages by transient interactions with an activator subunit. Despite their central role in ordering mitotic events, the mechanisms that govern APC/C-activator interactions remain poorly understood.

The APC/C is a ubiquitin ligase (E3) of the RING family, which catalyzes the assembly of polyubiquitin chains on lysine side chains within specific substrates, leading to their degradation by the proteasome. The APC/C is composed of 13 subunits that form a stable 1.3 MDa core enzyme^3,4^. Its activation requires association with an activator subunit, either Cdc20 or Cdh1, which acts as a substrate receptor and also stimulates enzymatic activity^3-5^. The two activators act in sequence to promote degradation of a series of substrates. From prometaphase to anaphase of mitosis, the APC/C associates with Cdc20 to ubiquitinate securin and mitotic cyclins, driving sister-chromatid separation and the completion of anaphase^6-8^. Cdc20 dissociates from the APC/C in late anaphase and is replaced by the second activator, Cdh1. Cdh1 maintains APC/C activity through late mitosis and during the subsequent G1, resulting in the sequential degradation of numerous Cdh1-specific substrates^9-11^. When the cell enters the next cell cycle in late G1, Cdh1 dissociation inactivates the APC/C, thereby allowing cyclins and other APC/C targets to accumulate. Thus, transient association of the APC/C with two different activator subunits organizes the timing of cell-cycle events.

Cdc20 and Cdh1 are structurally related proteins containing a seven-blade β-propeller WD40 domain that is the primary site of substrate binding^3,12^. This domain is anchored to the APC/C by sequences in flanking N- and C-terminal regions^3,5,13^. The N-terminal segment includes highly conserved interaction motifs, such as the C-box, that are critical for association with the APC/C core. Upon binding the APC/C, the unstructured regions around the C-box form a helical bundle that interacts extensively with a cavity in the APC/C^14^. Binding of activator to the APC/C also depends on a dipeptide IR motif at the activator C-terminus, which interacts with a binding groove on the Cdc27/Apc3 subunit of the APC/C^3,5,15^. Either of these N- or C-terminal motifs is indispensable for activator binding, and mutations in these regions abolish APC/C-activator interactions.

APC/C substrates interact with the activator WD40 domain via short linear sequence motifs called degrons, of which the D-box, KEN box and ABBA motif are the most common^12^. Cooperative binding of multiple degron sequences to the activator subunit provides the affinity required for efficient and processive ubiquitination, and the affinity of degron binding is likely to influence the timing of degradation in vivo^16-18^. Each of the three major degrons binds to a specific binding site on the activator WD40 domain. The KEN-box and ABBA motif interact directly with the WD40 domain, while the D-box interacts with both the WD40 domain and the adjacent core APC/C subunit Apc10/Doc1^3,4,13,19^. Bivalent D-box binding enhances APC/C-activator interactions by bridging the WD40 domain to the APC/C, providing additional interaction surfaces that stabilize the position of the activator on the APC/C^20-23^.

To gain more insight into the molecular mechanisms of APC/C-activator interactions, we analyzed the dissociation dynamics of activators *in vitro*. Using biochemical methods, we discovered that nucleic acids and other polyanions in cell lysates are able to rapidly dissociate activators from the APC/C. Interestingly, substrate D-box interactions between activator and the core APC/C reduce activator dissociation by polyanions, providing a mechanism that enhances activator binding when the enzyme is occupied with high-affinity substrate.

## RESULTS

### An activity in cell lysates dissociates Cdh1 and Cdc20 from the APC/C

The transient binding of activators to the APC/C during the cell cycle indicates that the activator dissociation rate in the cell must be sufficiently high to allow loss of activator from the APC/C over short time scales of minutes (Fig. 1a). To investigate the dynamics of activator binding, we developed an assay to measure activator dissociation from the budding yeast APC/C *in vitro* (Fig. 1b). We used translation *in vitro* to prepare ^35^S-labeled activator proteins, which were incubated with APC/C that had been immunopurified on magnetic beads from asynchronous cells carrying TAP-tagged APC/C subunit Cdc16. Following extensive washing and dilution, the amount of bound activator was then measured over time to estimate an activator dissociation rate (Fig. 1b).

**Figure 1.**
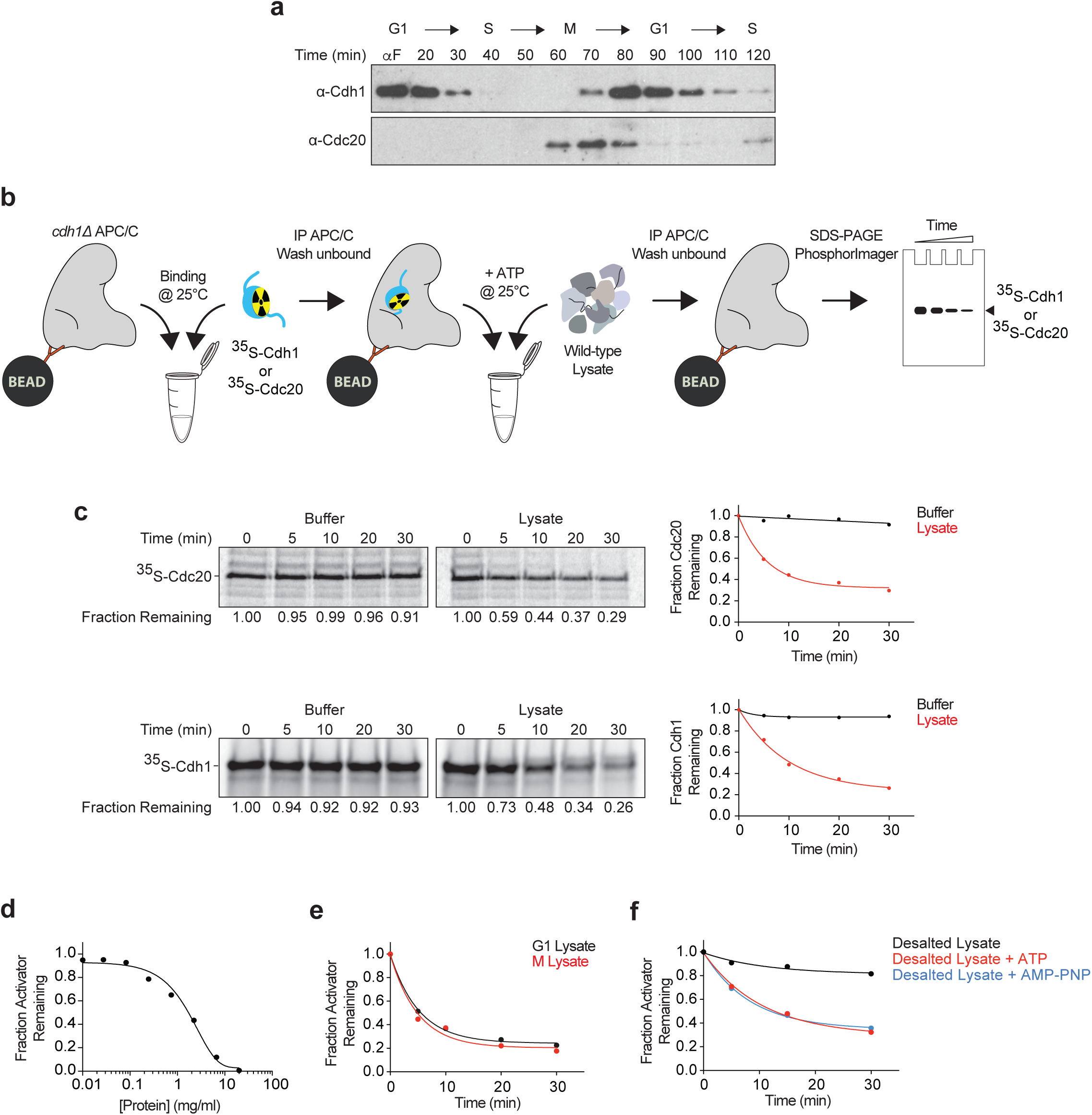
A biochemical activity in yeast cell lysates dissociates activators from the APC/C. **a.** Cells expressing TAP-tagged Cdc16 were arrested at G1 with 1 μg/ml *α*-factor for 3 h and released into YPD. APC/C was immunoprecipitated at the indicated time points and western blotted for Cdh1 (upper) and Cdc20 (lower). **b.** This schematic illustrates our APC/C-activator binding assay. *cdh1Δ* APC/C (TAP-tagged on the Cdc16 subunit) was immunopurified on magnetic beads and incubated with ^35^S-Cdc20 or ^35^S-Cdh1 produced by translation *in vitro*. After removing unbound activators by washing, buffer or yeast lysates (2.5 mg/ml protein concentration) was added in the presence of 5 mM ATP. At various times, APC/C was pulled down from the reaction mix and the amount of bound activator determined by SDS-PAGE and PhosphorImager analysis. Due to the extreme dilution of APC/C-activator complexes on the beads, rebinding of dissociated activator is expected to be negligible under these conditions. **c.** As described in panel **b**, we measured dissociation of radiolabeled Cdh1 (top) or Cdc20 (bottom) from immobilized APC/C over the indicated time course (at 25°C). Buffer control contains lysis buffer and 5 mM ATP. Fraction remaining was measured by calculating the ratio of activator signal at indicated time points to zero time point signal. Results were plotted and fitted to an exponential one phase decay equation in GraphPad Prism. **d.** Cdh1 dissociation reactions were performed as in panel **c** in the presence of serially diluted yeast lysate, supplemented with 5 mM ATP. Remaining activator after 45 min is plotted as a function of protein concentration. **e.** Lysates were prepared from wild-type cells arrested in G1 using *α*-factor (1 μg/ml) or in metaphase with nocodazole (15 μg/ml) and tested in Cdh1 dissociation assays in the presence of 5 mM ATP. **f.** Yeast lysates were subjected to gel filtration in lysis buffer to remove molecules smaller than ∼5 kDa. Desalted lysates were supplemented with buffer (black), 5 mM ATP (red) or 5 mM AMP-PNP (blue) prior to the Cdh1 dissociation reaction.

Cdh1 and Cdc20 both exhibited negligible dissociation in 30 minutes (Fig. 1c, Left panels), suggesting that activators bind the APC/C with high affinity. These results are consistent with the extensive binding interfaces seen in structural studies of APC/C-activator complexes^5,13-15^. However, they are not consistent with the rapid dissociation that occurs *in vivo*. We hypothesized that active mechanisms might exist in the cell to promote activator dissociation. Indeed, we found that incubation of APC/C^Cdh1^ or APC/C^Cdc20^ with whole lysates of budding yeast cells promoted activator dissociation (Fig. 1c, Right panels). This effect was not due to disassembly of the APC/C or proteolysis of the APC/C or activator (Supplementary Figs. 1a, b). Similar results were obtained with APC/C that was tagged at another APC/C subunit, Apc1, suggesting that activator dissociation is not due to the instability of the tagged Cdc16 subunit (Supplementary Fig. 1c). Dissociation rate depended on lysate concentration, and robust activator dissociation was achieved with cell lysates containing over 1 mg/ml of protein (Fig. 1d). Cdc20 and Cdh1 activate the APC/C at different stages of the cell cycle. To test whether dissociation activity is temporally regulated in a similar fashion, we compared lysates from cells arrested with mating pheromone (G1 arrest) or microtubule-depolymerizing drug nocodazole (metaphase arrest). Dissociation was promoted equally by these two lysates (Fig. 1e), suggesting that the dissociation activity does not change during the cell cycle.

ATP hydrolysis is the major energy source for protein remodelers, which are capable of pulling subunits of protein complexes apart. To investigate whether activator dissociation in cell lysates is ATP-dependent, we used gel filtration to remove molecules smaller than 5 kDa from cell lysates. Depleted lysates showed no significant dissociation activity, but activity was restored by addition of 5 mM ATP (Fig. 1f). ATP alone did not promote activator dissociation (Fig. 1c, Left panels). Addition of AMP-PNP also restored activity to depleted lysates (Fig. 1f), indicating that ATP hydrolysis is not required for the activity.

Stimulation of the dissociation activity by AMP-PNP ruled out canonical chaperone systems such as Hsp70, Hsp90 and Hsp104, which require ATP hydrolysis to carry out their functions. We also excluded these proteins by finding that dissociation activity was present in lysates from cells carrying mutant forms of these proteins (data not shown). The lack of a requirement for ATP hydrolysis also ruled out protein phosphorylation, which is known to influence activator binding to the APC/C^10,19,24,25^. We further narrowed the possible candidates by estimating the size of the activity by gel filtration chromatography. The activity eluted with a broad peak of 80-100 kDa (Supplementary Fig. 2a). This molecular size eliminated the possibility that activator dissociation was due to large molecular machines such as the proteasome or the chaperonin CCT, both of which are thought to influence activator binding to the APC/C^26-29^.

### Purification and identification of the activator dissociation activity

We attempted to isolate the dissociation activity from yeast lysates using a variety of chromatographic methods and other purification approaches. During the course of these studies, we discovered that the activity is resistant to boiling. Incubation of cell lysates at 95°C for 10 min led to precipitation of most proteins in the lysate, but the dissociation activity remained in solution (Fig. 2a). We then incorporated heat treatment as a step in an effective purification scheme, as follows (Fig. 2b). We applied cell lysate to a hydroxyapatite column, from which the activity could be eluted with 100 mM phosphate. The eluate was boiled, and the supernatant was dialyzed and re-applied to the hydroxyapatite column. The activity no longer bound to the column but was instead collected in the flow-through fraction (Fig. 2c). This method resulted in a high yield of activity and a high degree of purity. The flow-through fraction of this preparation was used for further characterization of the activity.

**Figure 2.**
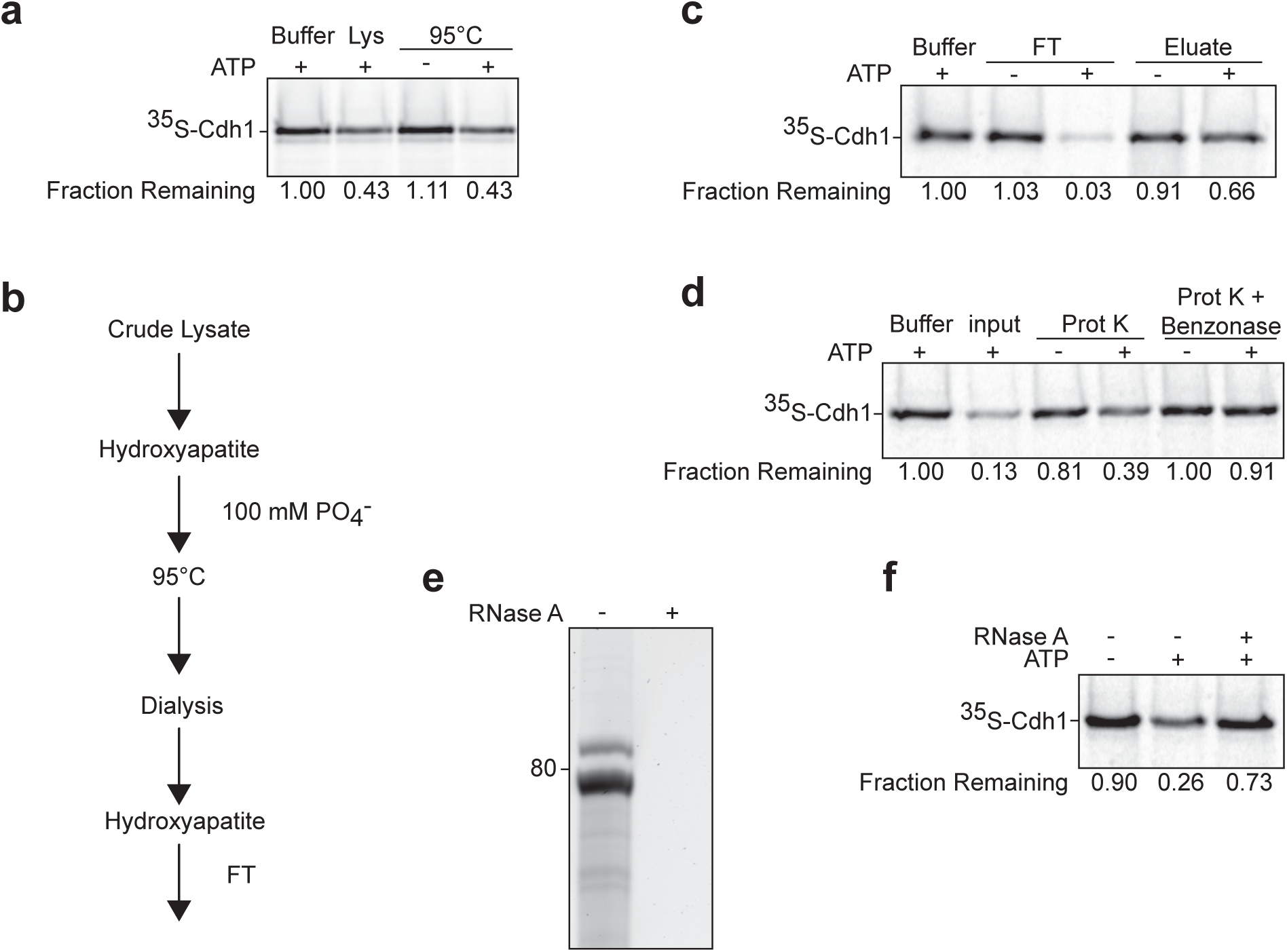
RNA in cell lysate is necessary and sufficient for activator dissociation. **a.** Cdh1 dissociation from APC/C was measured after 45 min in the presence of lysis buffer or yeast lysate, either untreated (Lys) or incubated at 95°C for 10 min. Prior to the experiment, boiled lysate was gel filtered and concentrated 5-fold. Dissociation reactions were performed with or without 5 mM ATP as indicated. **b.** Purification steps in the preparation of the dissociation activity used in panels **c**-**f**. Dissociation activity was found in the flow-through (FT) of the second hydroxyapatite column, which was also eluted with 100 mM phosphate. **c.** Flow-through (FT) and eluate fractions of the last hydroxyapatite step were tested in a Cdh1 dissociation reaction. Fractions were buffer-exchanged into reaction buffer containing 2.5 mM MgCl_2_. 5 mM ATP was added as indicated. The vast majority of the dissociation activity was found in the flow-through fraction. **d.** The hydroxyapatite flow-through fraction was incubated with buffer or Proteinase K (Prot K - 6 U/ml) at 37°C for 1 h. Proteinase K was heat inactivated at 85°C for 10 min and further inhibited with 1 mM PMSF. After Proteinase K treatment, samples were incubated with buffer or Benzonase (2000 U/ml) at 37°C for 1 h. Samples were supplemented with 5 mM ATP as indicated. **e.** Nucleic acid species in the flow-through fraction were extracted with phenol-chloroform, separated by a 10% TBE Urea polyacrylamide gel and stained with SYBR Safe. RNase A treatment (0.2 mg/ml) was performed at 37°C for 30 min (right lane). **f.** Phenol-chloroform-extracted RNA species from panel **e** were tested for Cdh1 dissociation with or without 3 mM ATP in the presence of a reaction buffer containing 2.5 mM MgCl_2_. RNase A treatment (0.2 mg/ml) was performed at 37°C for 30 min (right lane).

This fraction contained abundant activity despite having a very low protein concentration, leading us to question whether the dissociation activity is composed of protein or other macromolecules. Indeed, activity remained after treatment with the general protease Proteinase K (Fig. 2d), which led to the degradation of all detectable protein in the preparation (Supplementary Fig. 2b). However, treatment with a general nuclease, Benzonase, diminished the activity (Fig. 2d). Purification and analysis of the nucleic acids in the preparation revealed that the major nucleic acid species were about 80 nucleotides in length, and these species disappeared after RNase A treatment (Fig. 2e). We then purified the predominant RNA species and found that they removed Cdh1 from the APC/C in the presence of ATP, and RNase A treatment abolished the activity (Fig. 2f). These results argued that the RNA component is necessary and sufficient to dissociate activator subunit from the APC/C in an ATP-dependent manner.

### Nucleic acids and polyphosphate promote dissociation of activators from the APC/C

To further understand what types of RNA molecules provide the dissociation activity, we cloned and sequenced the major RNA species in our active preparation, revealing that the major RNAs in the preparation were tRNAs and rRNA fragments (see Methods). tRNA^Gln^ and tRNA^Ser^ were particularly abundant in the sample. To test their activity directly, tRNA^Gln^ and tRNA^Ser^ were transcribed *in vitro* and gel purified. Both species removed Cdh1 from the APC/C in the presence of ATP, with similar half-maximal dissociation occurring at concentrations of 0.9 μM and 0.7 μM, respectively (Fig. 3a).

**Figure 3.**
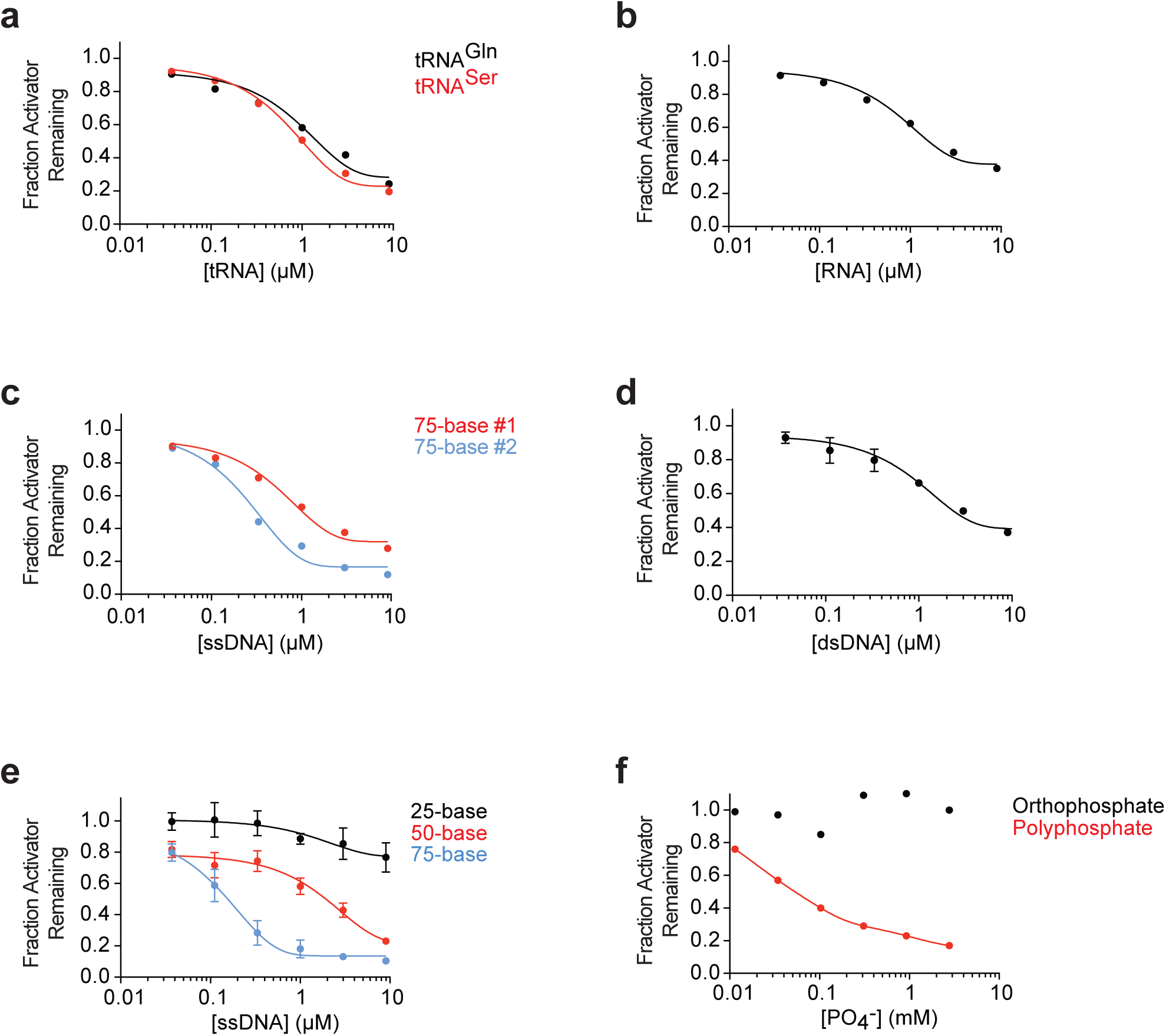
Nucleic acids and polyphosphate promote activator dissociation. **a.** tRNA sequences for glutamine (tRNA^Gln^) or serine (tRNA^Ser^) were transcribed *in vitro*, purified, and incubated in Cdh1 dissociation reactions for 45 min. Fraction of Cdh1 remaining at zero RNA concentration was not plotted on the log scale and taken as 1.0. **b-d.** Cdh1 dissociation reactions were performed in the presence of: **b.** a degenerate 74-nucleotide long RNA pool, transcribed *in vitro*; **c.** two random 75-base ssDNA oligos complementary to each other; and **d.** a double-stranded DNA formed by annealing the two single-stranded DNAs in panel **c**. Reactions were supplemented with 2.5 mM MgCl_2_ and 3 mM ATP. Error bars represent two independent experiments. **e.** Single-stranded DNA oligonucleotides of varying lengths were tested for Cdh1 dissociation in the presence of 2.5 mM MgCl_2_ and 3 mM ATP. Similar results were obtained in three independent experiments. **f.** Cdh1 dissociation reactions were performed at the indicated concentrations of orthophosphate (black) or polyphosphate (red) in a reaction buffer containing 3 mM ATP and no magnesium.

To determine if the activity was specific for tRNA or could be seen with any RNA species, we transcribed a pool of random RNA sequences of the same length as the glutamine tRNA used above. These RNA molecules removed activator from the APC/C at similar concentrations as tRNA (0.8 μM, Fig. 3b), suggesting that activator dissociation could be achieved by a variety of RNA sequences. To determine if this mechanism is a general function of nucleic acids, we synthesized two random, complementary 75-base single-stranded DNA oligonucleotides. Both sequences dissociated the activator subunit from the APC/C even more efficiently than RNA, with half-maximal concentrations of 0.2 μM and 0.6 μM (Fig. 3c). These results suggest that nucleic acids, independent of their sequence, are capable of promoting dissociation of activators from the APC/C.

The above experiments were performed with single-stranded nucleic acid polymers. Compared to double-stranded species, single-stranded nucleic acids exhibit more flexibility and heterogeneity in secondary structure. Slight variations in the dissociation activity of the sequences we tested (particularly differences between two complementary DNA oligonucleotides) suggest that nucleic acid structure and flexibility could play a role in disruption of APC/C-activator interactions. We explored this issue further by annealing the two complementary 75-nucleotide DNA oligonucleotides to reduce their flexibility and structural complexity. This double-stranded DNA dissociated activators less efficiently than either single strand (1 μM, Fig. 3d). Thus, efficient dissociation might be enhanced by the flexibility or secondary structures of single-stranded nucleic acids.

We also tested the effect of nucleic acid length on activator dissociation. A 75-base single-stranded DNA molecule was more than 5-fold (half maximal concentration 0.15 μM) more potent in activator dissociation than a 50-base oligonucleotide (1.8 μM). A 25-base oligonucleotide had little effect even at high concentrations (Fig. 3e). The higher activity of longer polymers could result from their greater flexibility and secondary structures. Activator dissociation might also depend on nucleic acid interactions with multiple distant sites on the APC/C and/or activator.

Given that both DNA and RNA can efficiently remove activators from the APC independent of nucleotide sequence, we next investigated if the negatively-charged phosphate backbone of nucleic acids is responsible for activator dissociation. We tested the activity of polyphosphate, long chains of phosphate molecules that are found at high concentrations in cells from every branch of life^30^. In addition to providing energy storage, polyphosphate is believed to alter protein structure and stability through extensive electrostatic interactions with protein surfaces^31^. We found that a mixed length polyphosphate preparation (45-160 phosphates per chain) was capable of dissociating activators, while similar concentrations of orthophosphate had no effect (Fig. 3f). Polyphosphate concentrations required for activator dissociation were higher than those observed with nucleic acids, even if nucleic acid concentration is expressed in terms of total phosphate concentration. We suspect that this reduced potency is due to heterogeneity in polyphosphate chain length. Taken together, our results show that biological polymers containing long chains of negatively-charged phosphates are potent catalysts of APC/C-activator dissociation.

### ATP promotes activator dissociation by sequestering magnesium ions

Our studies suggested that nucleic acids require ATP at millimolar concentrations for efficient activator dissociation (Fig. 2f, Supplementary Fig. 3a). Given that ATP alone, even at high concentrations, is not sufficient to dissociate activators (Supplementary Fig. 3a), and its hydrolysis is not necessary (Fig. 1e), it was likely that ATP had other roles in our reactions. To explore this issue further, we tested whether other nucleotides could substitute for ATP. The purified RNA preparation from the flow-through fraction of Fig. 2b displayed dissociation activity when supplemented with 3 mM ATP, AMP-PNP or GTP, and less effectively with ADP, but not with AMP (Fig. 4a). We hypothesized that dissociation might simply require molecules containing two adjacent phosphates. Consistent with this possibility, we found that inorganic pyrophosphate and tri-phosphate promoted the ability of DNA to dissociate the activator (Fig. 4b).

**Figure 4.**
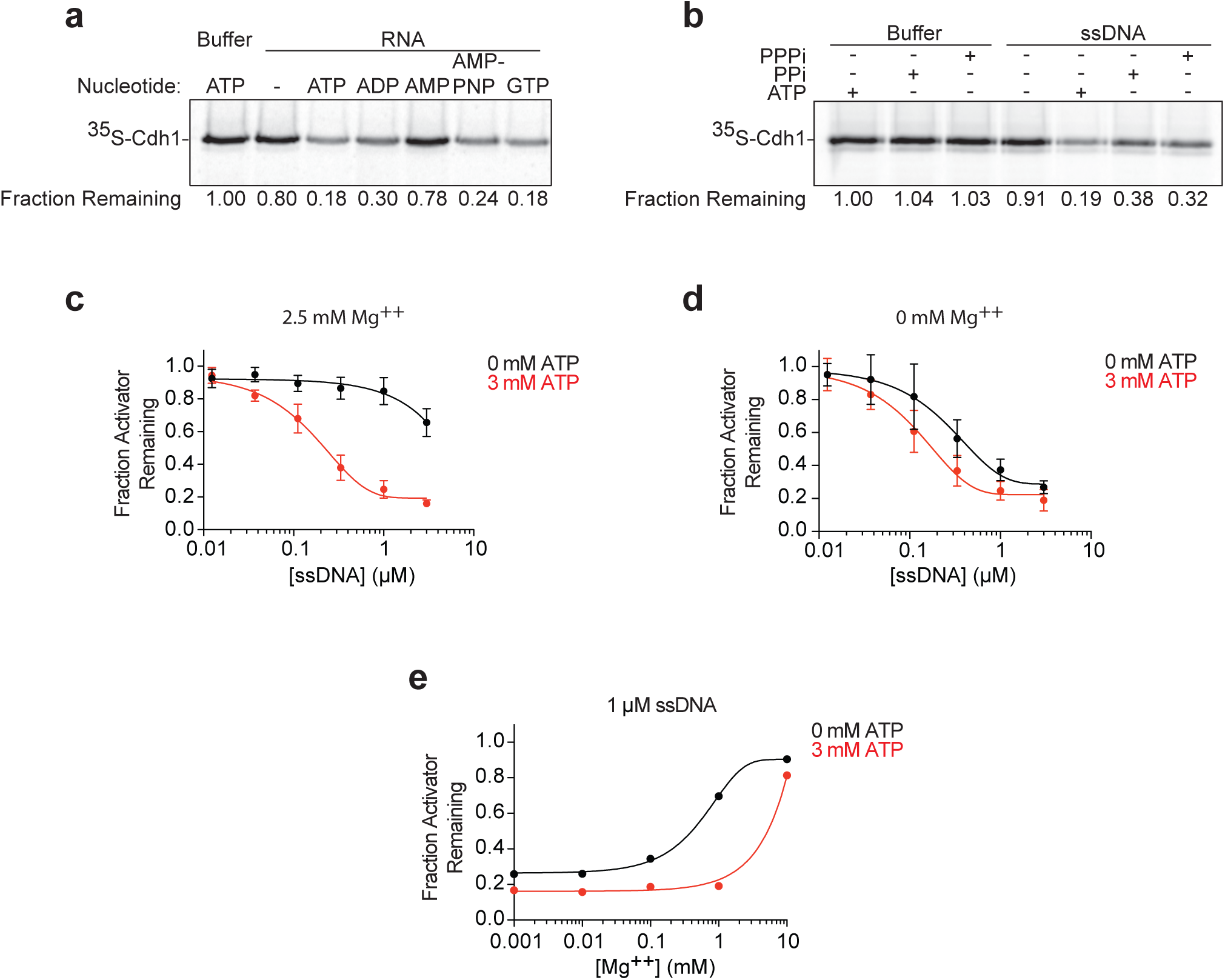
ATP promotes activator dissociation by sequestering magnesium ions. **a.** The effect of the indicated nucleotides on Cdh1 dissociation was tested using phenol-chloroform extracted RNA species from the hydroxyapatite flow-through fraction (Fig. 2b). Reactions were supplemented with 2.5 mM MgCl_2_ and 3 mM of the indicated nucleotides. **b.** 3 mM ATP, inorganic pyrophosphate (PPi), or tri-phosphate (PPPi) were tested in a Cdh1 dissociation assay with reaction buffer or 1 μM 75mer ssDNA oligonucleotide. Reactions contain 2.5 mM MgCl_2_. **c, d.** ssDNA-mediated activator dissociation requires ATP. A 75mer ssDNA oligonucleotide was tested in the Cdh1 dissociation assay in the presence (**c**) or absence (**d**) of 2.5 mM MgCl_2_. Reactions were supplemented with 0 mM (black) or 3 mM ATP (red). Error bars represent two independent experiments. **e.** The inhibitory effect of various MgCl_2_ concentrations was analyzed in Cdh1 dissociation reactions. Reactions were performed using 1 μM 75mer ssDNA with (red) or without (black) 3 mM ATP.

Phosphates on nucleic acids, polyphosphate chains and free ATP interact with divalent cations such as Mg^2+^. If polymeric negative charges are critical for the ability of nucleic acids and polyphosphate to dissociate activators, then magnesium ions could neutralize this charge and thereby block activator dissociation. Our dissociation reactions contain 2.5 mM Mg^2+^ ions, and we therefore hypothesized that millimolar concentrations of ATP promote activator dissociation by sequestering these ions. Consistent with this hypothesis, magnesium ions inhibited DNA- or polyphosphate-dependent activator dissociation in the absence of ATP (Fig. 4c, d; Supplementary Fig. 3b). ATP was not required when single-stranded DNA was used in the absence of magnesium, and the reaction was only slightly enhanced when supplemented with ATP (Fig 4d). In reactions containing both ATP and magnesium, we observed that the inhibitory effect of magnesium occurred only when it was in molar excess of ATP (Fig. 4e, Supplementary Fig. 3c). These findings indicate that ATP sequesters magnesium ions to enhance the negative charge of phosphate-containing polymers, which is required for activator dissociation.

### Polyanions enhance APC/C substrate selectivity

The APC/C ubiquitinates its targets only when bound by an activator, so dissociation of activators by polyanions should reduce APC/C activity if polyanions are capable of dissociating functional activators. We tested this possibility by measuring APC/C ubiquitination activity toward radiolabeled Pds1/securin *in vitro.* As predicted, pre-incubation of APC/C^Cdh1^ with single-stranded DNA prior to the ubiquitination reaction resulted in lower Pds1 ubiquitination (Fig. 5a).

**Figure 5.**
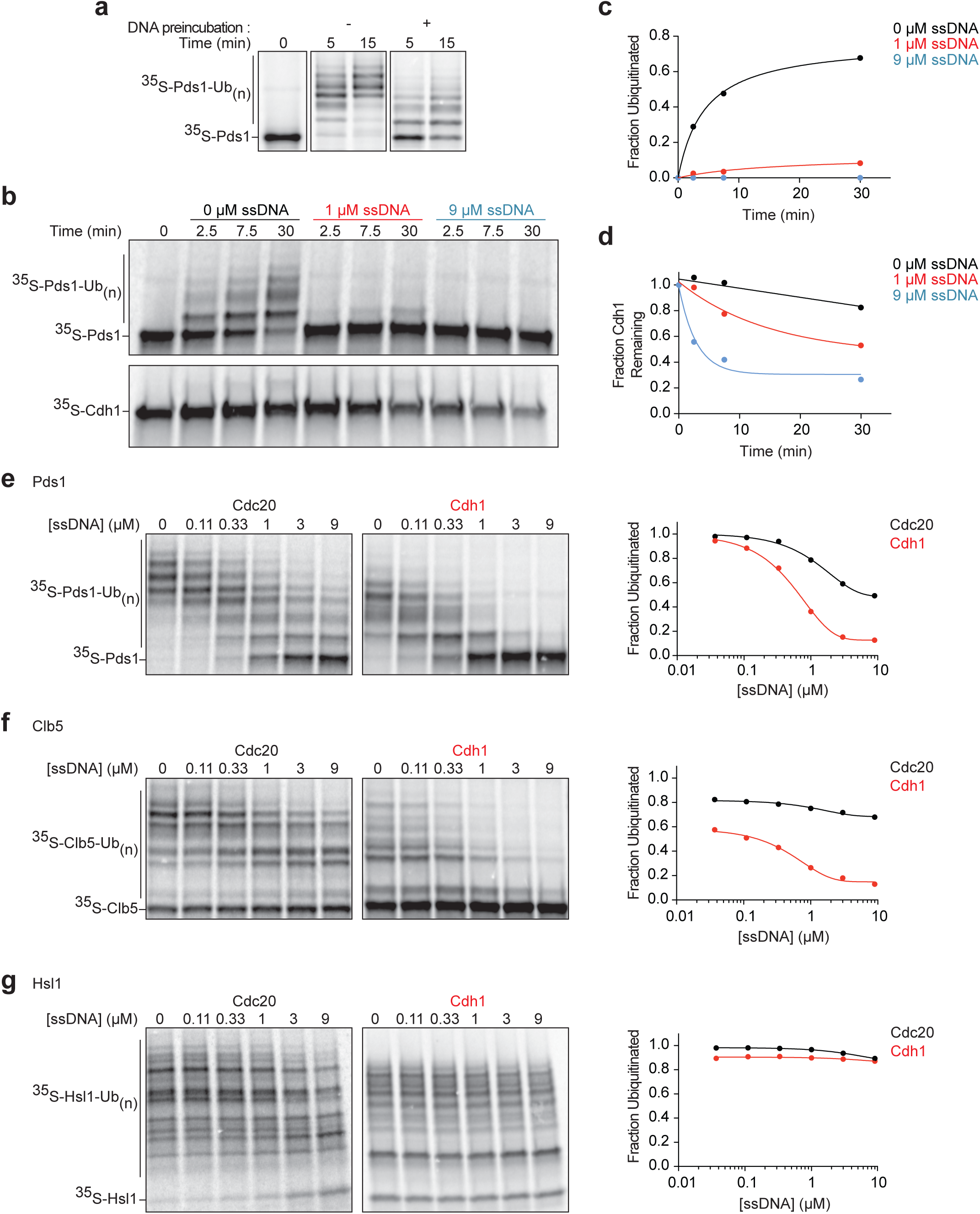
Polyanions modulate APC/C activity toward different substrates. **a.** APC/C was immunopurified on beads and incubated with unlabeled Cdh1 (translated *in vitro*). Following washing, APC/C^Cdh1^ was incubated 30 min at 25°C with buffer (-) or 1 μM 75mer single-stranded DNA (+) in the presence of 3 mM ATP. After 2 wash steps, ^35^S-Pds1 substrate was added to the APC^Cdh1^ and ubiquitination reactions were performed at 25°C with methylated ubiquitin and Ubc4 as the E2. Reactions were terminated at the indicated times and analyzed by SDS-PAGE and PhosphorImager. **b.** APC/C was purified from cells carrying Apc1-GFP using anti-GFP on magnetic beads. Ubiquitination reactions were performed in two parallel experiments using ^35^S-Pds1 and unlabeled Cdh1 to monitor Pds1 ubiquitination (upper panel) or ^35^S-Cdh1 and unlabeled Pds1 to detect activator dissociation (lower panel). 75mer ssDNA was added to the reactions at the indicated concentrations, with 3 mM ATP. Ubiquitination reactions were terminated at the indicated times and analyzed by SDS-PAGE and PhosphorImager. For the ^35^S-Cdh1 dissociation reactions, immobilized APC/C was washed prior to analysis. **c, d.** Quantification of results in panel **b**: **c.** Fraction of ubiquitinated ^35^S-Pds1 was measured by calculating the ratio of ubiquitinated substrate to total substrate in the reaction. **d.** ^35^S-Cdh1 dissociation was calculated by measuring ^35^S-Cdh1 that remained bound to the APC/C as in previous experiments. **e-g.** Immunopurified APC/C was pre-bound with unlabeled Cdc20 (left) or Cdh1 (right) and then used in ubiquitination reactions with one of three substrates (**e.** ^35^S-Pds1**; f.** ^35^S-Clb5; **g.** ^35^S-Hsl1(aa 667-872)). Reactions were performed in the absence or presence of the indicated concentrations of the 75mer ssDNA, plus 2.5 mM MgCl_2_ and 3 mM ATP. Ubiquitination reactions were performed at 25°C for 10 min (Hsl1) or 30 min (Pds1 and Clb5). Quantification of substrate ubiquitination is provided in the panels at right.

To better understand APC/C function in the presence of DNA, we simultaneously measured Pds1 ubiquitination and activator dissociation at various DNA concentrations. Even after just 2.5 minutes of incubation with DNA, Pds1 ubiquitination by APC/C^Cdh1^ was almost completely inhibited at DNA concentrations of 1 μM or higher (Fig. 5b, c), despite the fact that a significant amount of Cdh1 remained bound to the APC/C (Fig. 5b, d). After 30 minutes, ubiquitination activity remained inhibited and was accompanied by more Cdh1 dissociation (Fig. 5c, d). These results revealed that inhibition of APC/C activity by polyanions is not due simply to activator dissociation. Instead, they suggest that inhibition occurs in two steps: immediate inhibition of substrate ubiquitination followed by secondary inactivation due to activator dissociation.

We next analyzed the effect of DNA on the ubiquitination of a selection of substrates, using Cdh1 or Cdc20 as activator. We tested three substrates: Pds1 and S-phase cyclin Clb5, both well-established substrates of APC/C^Cdc20^ in metaphase; and Hsl1, a protein that is ubiquitinated primarily by APC/C^Cdh1^ in late mitosis but is also an excellent substrate for APC/C^Cdc20^ *in vitro*^14,20^. As before, we observed that APC/C^Cdh1^ activity toward Pds1 was inhibited by single-stranded DNA, with a half-maximal DNA concentration of 0.2 μM (Fig. 5e). Interestingly, Pds1 ubiquitination by APC/C^Cdc20^ was relatively resistant to inhibition (Fig. 5e). Results with the S-phase cyclin Clb5 displayed a similar trend, with greater inhibition of activity with APC/C^Cdh1^, and resistance to DNA with APC/C^Cdc20^ (Fig. 5f). On the other hand, DNA had little effect on ubiquitination of Hsl1 by APC/C with either activator (Fig. 5g).

The three substrates we tested are likely to have different affinities for different APC/C-activator complexes. Substrate affinity is reflected in the processivity of the ubiquitination reaction: a slower dissociation rate (and thus higher affinity) generally results in a greater number of ubiquitins attached during a single binding event^21,32,33^. Pds1 and Clb5 are both more processively modified by APC/C^Cdc20^ than APC/C^Cdh1^ (Lu et al., 2014) (Fig. 5e, f), suggesting that they have greater affinity for Cdc20 than Cdh1. Hsl1 is an unusual substrate that is modified with extremely high processivity with either activator (Fig. 5g), and it is known to contain an exceptionally high-affinity D-box that has been used extensively in structural studies of the APC/C^14^. Together, these results suggest that DNA is a less effective inhibitor of reactions with substrates that bind with high affinity, indicating that DNA and substrate compete, directly or indirectly, for binding to the APC/C-activator complex.

### D-box binding inhibits polyanion-mediated activator dissociation

Our evidence that high-affinity substrate ubiquitination is resistant to inhibition by DNA led us to test the possibility that high-affinity substrate binding also blocks the effect of DNA on activator dissociation. We purified a fragment of Hsl1 (aa 667-872) that contains the D-box and the KEN box and tested the effect of Hsl1 binding on single-stranded DNA-mediated dissociation. Hsl1 stabilized the bound activator and completely blocked DNA-driven dissociation of Cdh1 from the APC/C (Fig. 6a).

**Figure 6.**
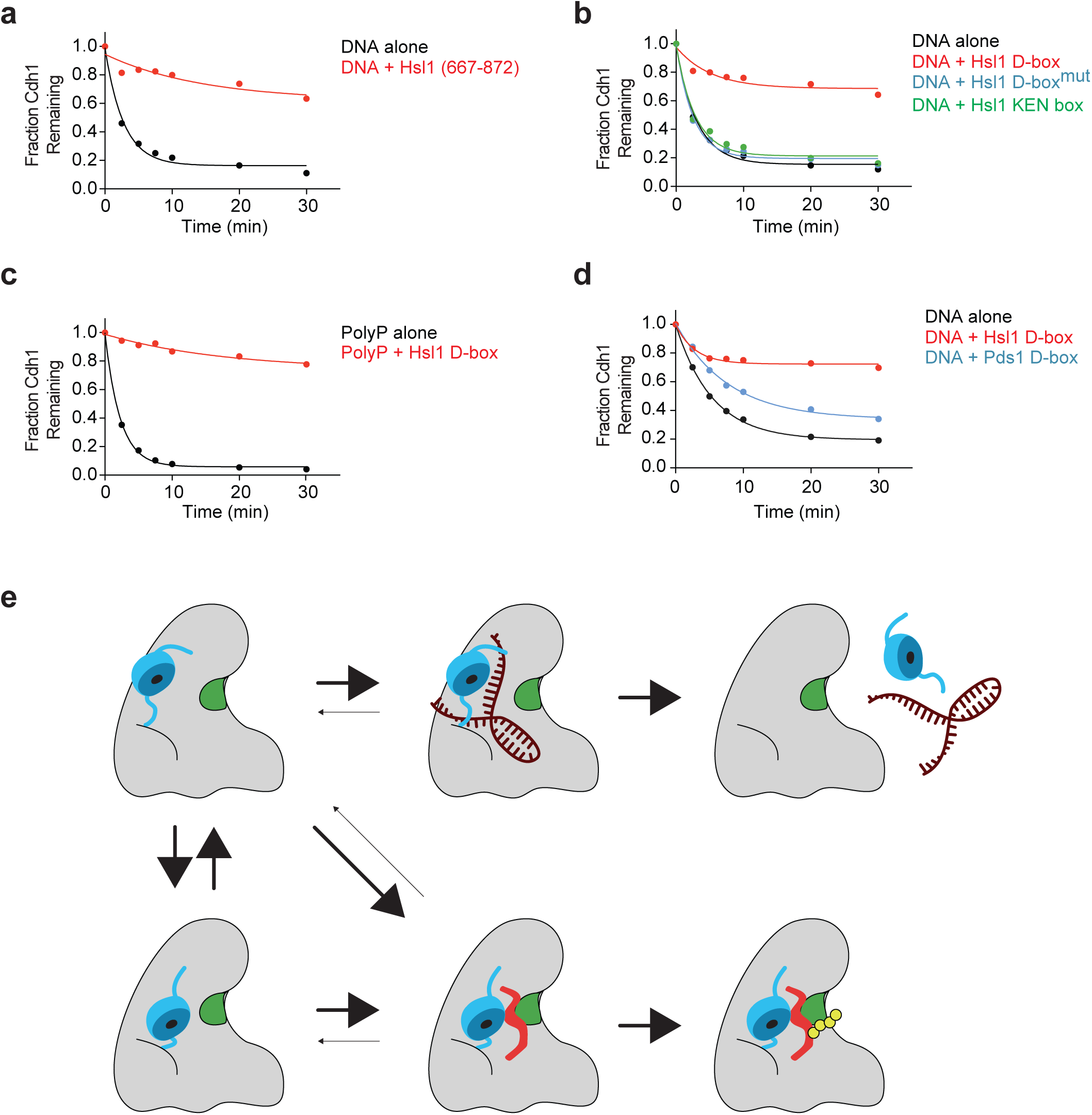
D-box-dependent substrate binding protects activator from dissociation. **a.** Cdh1 dissociation was measured with 1.5 μM 75-base ssDNA and 3 mM ATP, in the absence (black) or presence (red) of 4 μM purified Hsl1 fragment (aa 667-872). **b.** Cdh1 dissociation was measured in the presence of buffer (black), 4 μM Hsl1 D-box peptide (red), Hsl1 D-box peptide mutant (blue) or Hsl1 KEN box peptide (green). Reactions were performed in the presence of 1.5 μM 75mer ssDNA and 3 mM ATP. **c.** Cdh1 dissociation was measured in the presence of polyphosphate (1 mM total PO_4_ ^-^) with (red) or without (black) 4 μM Hsl1 D-box peptide. **d.** Cdh1 dissociation was measured in the absence (black) or presence of D-box peptides (4 μM) from Hsl1 (red) or Pds1 (blue). **e.** Speculative model for dynamic regulation of APC/C activity by polyanions. Activator binding to the APC/C is mediated by binding motifs located on flexible N- and C-termini. In the absence of substrate (left column), the activator WD40 domain (blue) is mobile, resulting in variable positioning of the WD40 domain relative to the core subunit Apc10 (green). We speculate that **a.** movement of the WD40 domain away from Apc10 (top row) exposes binding sites for polyanions such as polyphosphate or nucleic acids (brown), preventing substrate binding and thereby preventing WD40 interaction with Apc10. Polyanions thereby block the ubiquitination reaction and also promote activator dissociation, perhaps by interfering with interactions between APC/C and activator termini. When substrates are present (lower right), the D-box (red) links activator and Apc10, thereby enhancing activator binding and preventing polyanion-mediated inhibition. Processive substrate ubiquitination (yellow) is then possible.

The D-box of APC/C substrates is known to link the activator subunit to the core APC/C subunit Apc10, and previous work has shown that D-box binding enhances activator binding to the APC/C^20,21^. Similarly, we found that a peptide containing only the Hsl1 D-box was able to block the effects of DNA and polyphosphate on activator dissociation (Fig. 6b, c). A mutant D-box peptide, or a KEN box peptide, had no effect (Fig. 6b).

Polyanionic inhibition of APC^Cdh1^ activity toward Pds1 or Clb5 (Fig. 5e, f) might result because their D-boxes bind to Cdh1 weakly and cannot compete effectively with polyanions. Consistent with this idea, we found that a peptide containing the Pds1 D-box only partially blocked the ability of DNA to promote Cdh1 dissociation (Fig. 6d). In contrast, a Pds1 fragment (1-110aa) that contains the D-box greatly reduced the dissociation of Cdc20 (Supplementary Fig. 4), consistent with the notion that the inhibitory effects of DNA are blocked most effectively by high-affinity D-box binding. Together with our studies of ubiquitination reactions in Fig. 5, these results argue that a high-affinity D-box interaction holds the activator in a position that blocks the effects of polyanions on substrate ubiquitination and activator dissociation.

## DISCUSSION

Precise temporal control of APC/C activity during the cell cycle relies on short-lived interactions between the APC/C and the activator subunit. Our studies suggest that the transient binding of activator to the APC/C is unlikely to result from spontaneous dissociation, which is negligible *in vitro*. Instead, activator dissociation is accelerated by abundant cellular polyanions such as DNA, RNA and polyphosphate, which also inhibit APC/C activity. The inhibitory effects of polyanions are blocked by strong substrate binding, providing a mechanism to selectively enhance ubiquitination of high-affinity substrates.

There is growing evidence that negatively-charged biopolymers influence the structural dynamics of proteins in the cell. Nucleic acids and polyphosphate exhibit chaperone-like properties, accelerate protein folding, and serve as anti-aggregation agents by solubilizing a variety of protein aggregates *in vitro*^34-36^. Long stretches of phosphate residues also inhibit polymerization of multimeric proteins and stabilize soluble structures^37,38^. Our work reveals that polyanions also influence the dynamics of APC/C-activator interactions, specifically disrupting APC/C-activator binding while leaving the multi-subunit APC/C core intact.

We found that submicromolar concentrations of RNA and DNA promote APC/C-activator dissociation, while polyphosphate acts at submillimolar concentrations. These concentrations are far lower than RNA and polyphosphate concentrations inside the cell. tRNA alone can be found in cells at concentration of approximately 200 μM, while polyphosphate concentration is estimated to be as high as 100 mM^30,39^. Even if these molecules are partially neutralized by positively-charged ions such as Mg^2+^ and other charged macromolecules in the cell, it seems likely that there are sufficient quantities of various negatively-charged polymers to exceed the low concentrations needed *in vitro* to promote APC/C inhibition and activator dissociation.

The binding of activator to the APC/C is known to be regulated by multiple mechanisms. The best understood is phosphorylation of Cdh1 by cyclin-dependent kinases (Cdks), which is known to reduce Cdh1 binding to APC/C in the cell^10,24,40^. Several Cdh1 phosphorylation sites are found in N-terminal regions that mediate APC/C binding. These regions become less accessible to Cdks upon APC/C binding, suggesting that phosphorylation is likely to block association of free Cdh1 rather than promoting dissociation. Cdc20 binding to the APC/C, on the other hand, is controlled in part by its ubiquitination. Cdc20 levels in late mitosis decrease as a result of APC/C^Cdh1^-mediated degradation as well as autoubiquitination. Cdc20 autoubiquitination is also involved in disruption of the mitotic checkpoint complex that blocks APC/C^Cdc20^ activity during activation of the spindle assembly checkpoint^15,29^. However, there is no indication that activator phosphorylation or ubiquitination is required for the polyanion-mediated activator dissociation we observed in cell lysates or with purified components.

In addition to providing new insights into the control of APC/C-activator interactions, we uncovered new clues about APC/C-substrate binding. APC/C substrates contain combinations of one to three degrons, of which the D-box is particularly critical because of its role in linking the activator to the APC/C core. The amino acid sequences of the D-box and other degrons vary in different substrates, and it is likely that these sequences evolved to be accommodated by different APC/C-activator complexes. Substrate affinity for each of the two activators determines the substrate residence time. As a result, these variations are likely to influence processivity of ubiquitination and the timing of substrate degradation in vivo^17^. We found that substrate affinity also controls the ability of polyanions to inhibit APC/C activity: the binding of high-affinity substrates results in resistance to the effects of polyanions, while low-affinity substrates are less resistant. Thus, differences in enzyme processivity with substrates of different affinities are amplified in the presence of polyanions, providing a selectivity filter that promotes ubiquitination of high-affinity targets while suppressing background activity toward low-affinity substrates. Furthermore, the ability of substrate binding to stabilize activator binding could enhance APC/C-activator function at cell-cycle stages when abundant substrates are present.

Our observations with different substrates also suggest that polyanions compete with the substrate D-box for binding to the APC/C-activator complex. This competition provides clues about the mechanism of polyanion action. In the absence of substrate, activators are attached to the APC/C via flexible N- and C-termini, such that the globular WD40 domain does not stably interact with APC/C core and remains mobile. It is plausible that polyanions interact with the APC/C at sites adjacent to the activator that are exposed in the absence of a substrate (Fig. 6e, upper row); binding at these sites could prevent substrate binding (thereby inhibiting ubiquitination) while at the same time promoting activator dissociation by disrupting binding sites for one or both activator termini. In this model, when high-affinity D-box binding links the WD40 domain to Apc10, polyanions lose access to their interaction sites and have less impact on activator binding. Stabilization of activator, as a result, stimulates the catalysis of ubiquitin transfer (Fig. 6e, lower row).

With combined concentrations exceeding tens of millimolar, nucleic acids, polyphosphate chains, ATP and other charged macromolecules are in frequent close contact with proteins inside the cell. Transient interactions between polyanions and protein surfaces may create further crowding in the molecular microenvironment around proteins, while also altering the structural and functional dynamics of protein complexes and other macromolecules. Emerging roles of ATP beyond energy provision suggest that the electrostatic microenvironment around biomolecules has an impact on biochemical outputs^41^. Our understanding of polyanion biology is limited, but further studies could help us discover and characterize other examples of non-canonical molecular interactions that modulate complex enzyme functions.

## MATERIALS AND METHODS

### Synchronization and Western Blotting

All yeast strains were derivatives of W303. For western blotting of activators during the cell cycle, yeast cells carrying TAP-tagged Cdc16 (Strain yAM4: Cdc16-TAP::KanMX, MATa) were grown overnight and diluted the next morning. When cells reached OD 0.3, cells were synchronized at 30°C for 3 h with 1 μg/ml *α*-factor. Following release from the arrest by washing, cells were harvested at various times and lysed by bead beating in lysis buffer as described in the next section. TAP-tagged APC/C was immunoprecipitated with magnetic IgG beads. Cdc20 and Cdh1 were subjected to western blotting with polyclonal goat yC-20 (sc-6731) and yC-16 (sc-8959), respectively.

### Activator binding and dissociation

A yeast strain carrying Cdc16-TAP and lacking Cdh1 (Strain DOM1226: cdh1::LEU2, Cdc16-TAP:HIS, MATa) ^22,28^ was harvested at OD_600_=1 and flash frozen in liquid nitrogen. Cell pellets were lysed by bead beating in lysis buffer (25 mM HEPES pH 7.6, 150 mM KOAc, 2.5 mM MgCl_2_, 10% Glycerol, 0.5% Triton X-100, 1 mM PMSF, 1 mM DTT and EDTA-free protease inhibitor mix) and centrifuged at 20,000xg for 15 min at 4°C. The APC/C was immunoprecipitated with IgG-coupled magnetic beads (Invitrogen #14301) at 4°C. Cdc20 and Cdh1 were translated *in vitro* as described^21,22,28^, using the TnT Quick-Coupled reticulocyte lysate system (Promega #L1170) and ^35^S-Methionine (Perkin Elmer #NEG709A001MC). APC/C on beads was incubated with reticulocyte lysate containing Cdc20 or Cdh1 for 30 min at 25°C, followed by washing three times with lysis buffer to remove unbound activator. Dissociation reactions were performed at 25°C by incubating APC/C^Cdh1^ or APC/C^Cdc20^ with lysis buffer or with wild-type lysates prepared by bead beating in lysis buffer as above and supplemented with 5 mM ATP. Nucleic acid-dependent dissociation assays were performed with 3 mM ATP unless otherwise noted. To terminate the reactions, APC/C was washed with the same buffer and separated by 10% SDS-PAGE. Gels were dried on Whatman paper, exposed to a Phosphor Screen and scanned in a Typhoon 9400 Imager. Images were analyzed using ImageQuant (GE Healthcare). Fraction of Cdc20 and Cdh1 remaining on the APC/C was calculated relative to the zero timepoint or buffer control. In Fig. 5b, APC/C was immunopurified from lysates of a strain lacking Cdh1 and carrying GFP-tagged Apc1 and TAP tagged Cdc16 (Strain NYH14: cdh1::LEU2, Cdc16-TAP:HIS, Apc1-eGFP:CaURA3, MATa). The Hsl1 fragment used in Fig. 6a was expressed in insect cells and purified using a Strep-tag purification column. Degron peptides used in Fig. 6b-d were purchased from CPC Scientific. Peptide sequences are listed in Supplementary Table 1.

### Partial purification of dissociation activity

Wild-type yeast cells were collected at OD_600_=1.0, washed twice with PBS, and resuspended in water and flash frozen in liquid nitrogen. Cells were lysed using a coffee grinder in a buffer containing 25 mM HEPES pH 7.6, 25 mM KOAc, 2.5 mM MgCl_2_, 10% Glycerol, 0.5% Triton-X 100, 1 mM PMSF and protease inhibitor mix. Lysate was cleared by centrifugation at 100,000xg for 1 h and filtered through a 0.22 μm cellulose filter. Cleared lysate was applied to a Bio-Rad CHT type II ceramic hydroxyapatite column using a peristaltic pump. The column was washed with 15 column volumes of buffer HA-A (25 mM HEPES 7.6, 25 mM KOAc, 10% Glycerol) containing 2.5 mM MgCl_2_. Samples were eluted with a step phosphate gradient using seven column volumes of the same buffer supplemented with 50 mM, 100 mM, 250 mM, or 500 mM PO4 ^-^. Small portions of the flow-through and elution fractions were concentrated, buffer-exchanged into Buffer HA-A with 2.5 mM MgCl_2_ and tested for activator dissociation in the presence of 3 mM ATP. The activity eluted at 50 mM and 100 mM PO_4_ ^-^ concentrations. The eluted activity was incubated at 95°C for 15 min and centrifuged at 100,000xg for 1 h at 4°C to pellet precipitated molecules. Cleared boiled samples were dialyzed into Buffer HA-A with 2.5 mM MgCl_2_ overnight using SnakeSkin Dialysis Tubing (Thermo Fisher #68035) at 4°C. Dialyzed samples were re-applied to the hydroxyapatite column. The vast majority of the dissociation activity did not bind to the column and was collected in the flow-through fraction. This fraction was dialyzed into Buffer HA-A overnight and concentrated 10-fold for further characterization in dissociation assays. The activity in this fraction was stable for more than 2 weeks at 4°C with no significant loss of activity.

### RNA preparation and sequencing

RNA species in the flow-through fraction were separated by 10% TBE-urea polyacrylamide gel, and major RNA species (∼80 nucleotides) were extracted from the gel and ethanol precipitated. A 3’ DNA adapter (CTATAGTGTCACCTAAATTAATACGACTCACTATAGGG) that contains 5’ phosphate and 3’ spacers was first 5’-adenylated using a 5’-adenylation kit (NEB #E2610-S) at 65°C for 1 h, then ligated to purified RNA species using T4 RNA Ligase 2 Truncated (NEB #M0242S) at 25°C for 1 h. The ligation reaction was separated on a 10% TBE-urea polyacrylamide gel, and ligated products were gel purified. Next, a 5’ RNA adapter (GCAATTAACCCTCACTAAAGGAGTCGT) lacking 5’ phosphate was ligated with T4 RNA Ligase 1 (NEB #M0204S). Ligation products were gel-purified, and cDNA was synthesized using RT primer (CCCTATAGTGAGTCGTATTAATTTAGGTGACACTATAG) and Superscript IV reverse transcriptase (Thermo-Fisher #18090010). After RNase H digestion, cDNAs were amplified using T7 (TAATACGACTCACTATAGGG) and T3 (GCAATTAACCCTCACTAAAGG) primers. PCR products were cloned into TOPO vectors and transformed into DH5-*α* cells. After overnight growth and plasmid isolation, sequencing was performed using M13 Forward and Reverse primers. Alignment of sequencing results with the *S. cerevisiae* S288C genome revealed the presence of tRNAs (tQ(UUG), tS(AGA), tG(CCC), tD(GUC), tE(CUC), tW(CCA), tR(CUC), tL(UAG)) and similar lengths of rRNA fragments (RDN25-1).

### Nucleic acid preparation for dissociation reactions

Sequences used in dissociation assays are listed in Supplementary Table 1. RNA sequences were transcribed *in vitro* as previously described^42^ using T7 RNA Polymerase (NEB #M0251). Chemically synthesized DNA oligonucleotides or transcribed RNA species were separated on a TBE-urea polyacrylamide gel, purified, ethanol precipitated and resuspended in Buffer HA-A with or without 2.5 mM MgCl_2_. Medium chain polyphosphate was obtained from Kerafast (p100 #EUI005) and resuspended in Buffer HA-A without MgCl_2_.

### APC/C ubiquitination assay

Yeast cells carrying Cdc16-TAP and lacking Cdh1 were collected at OD_600_=1, flash frozen, and lysed using beat beating in lysis buffer. APC/C was immunopurified on IgG beads and activated by *in vitro* translated Cdc20 or Cdh1. Cdc20 and Cdh1 used in Fig. 5e-g were expressed in insect cells using the baculovirus system and purified using Strep-tag purification methods. For APC/C substrates, Pds1, Hsl1 fragment (aa 667-872), and Clb5 were C-terminally ZZ-tagged, cloned into plasmids under control of the T7 promoter, and translated *in vitro* using ^35^S-Methionine. Substrates were purified using IgG-coupled magnetic beads and cleaved with TEV protease (Thermo Fisher #12575015). E1 and E2 (Ubc4) were expressed in *E. coli* and purified as previously described^43,44^. E2 charging was performed in a reaction containing 0.2 mg/ml Uba1, 2 mg/ml Ubc4, 2 mg/ml Methylated ubiquitin (Boston Biochem #U-501) and 1 mM ATP at 37°C for 30 minutes. The ubiquitination reaction was initiated by mixing activated APC/C, E2-ubiquitin conjugates, purified substrate, and buffer or indicated concentrations of single-stranded DNA at 25°C. Reactions were terminated by addition of 2X SDS Sample Loading Dye and separation by SDS-PAGE.

## ACKNOWLEDGEMENTS

We thank Robert Cohen, David Agard, Peter Walter, Mark Von Zastrow and members of the Morgan lab for their technical and intellectual insights; Nairi Hartooni for providing purified Cdc20, Cdh1 and Hsl1 fragment; Kelsey Hickey for discussions and comments on the manuscript. This work was supported by the National Institute of General Medical Sciences (R35-GM118053, to DOM) and an HHMI International Student Research Fellowship (AM).

## AUTHOR CONTRIBUTIONS

A.M. conceived the project, performed the experiments, and wrote the paper, with guidance from D.O.M.

## Supplementary Material

**Supplementary Figure 1.**
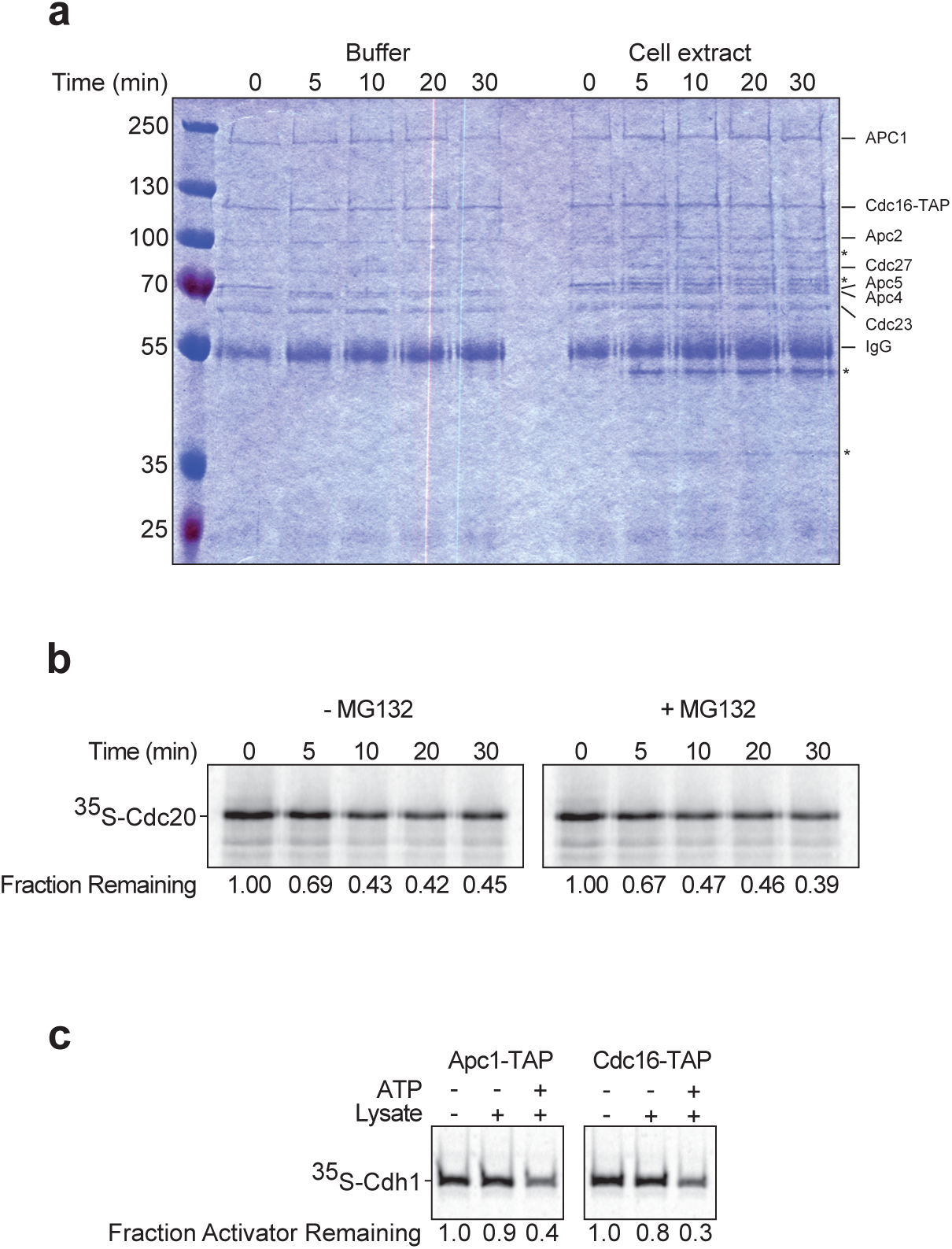
Characterization of dissociation activity. **a.** APC/C subunits remain intact during the activator dissociation reaction. As in Fig. 1c, dissociation reactions were performed with buffer or yeast lysates, terminated at the indicated times, and analyzed by SDS-PAGE and staining with Coomassie Blue. Nonspecific lysate proteins that bind to IgG beads are indicated by asterisks. **b.** Cdc20 dissociation reactions were performed with yeast lysate (2.5 mg/ml) in the absence (left) or presence (right) of the proteasome inhibitor MG132 (50 μM) plus 5 mM ATP. **c.** APC/C was immunopurified from lysates of cells carrying TAP-tagged Apc1 (left) or Cdc16 (right), and Cdh1 dissociation reactions were performed with yeast lysate plus 5 mM ATP.

**Supplementary Figure 2.**
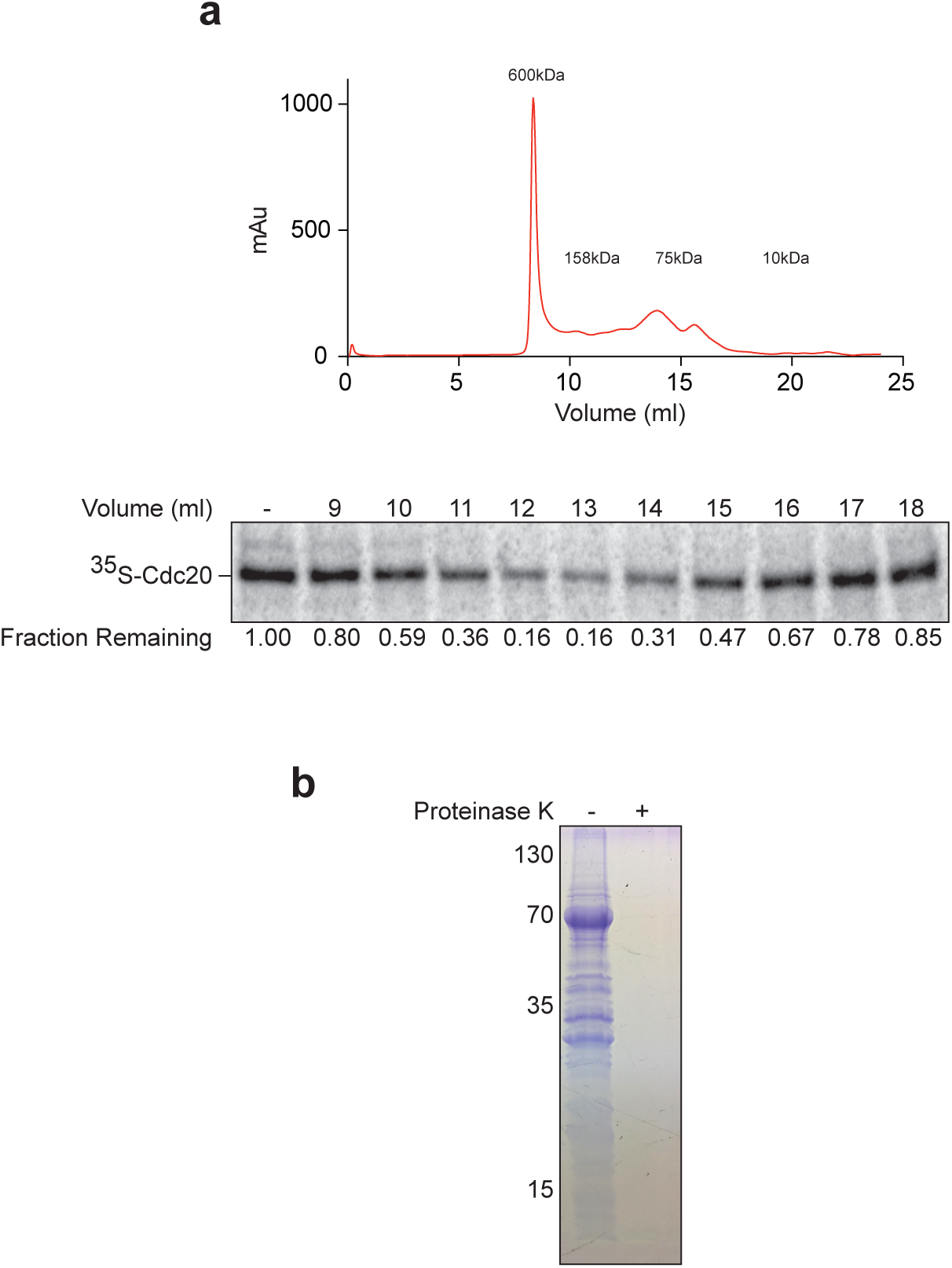
Molecular size of dissociation activity. **a.** Yeast lysates were applied to a DEAE-Fast flow ion exchange column, and dissociation activity was eluted with 500 mM KOAc, concentrated and applied to a Superdex 200 gel filtration column. Top panel shows the UV absorbance (280 nm) of the eluate. In the bottom panel, fractions were concentrated 10-fold and used in a Cdc20 dissociation reaction, supplemented with 5 mM ATP. **b.** As described in Fig. 2d, the hydroxyapatite flow-through fraction from Fig. 2b was treated with buffer (left) or 0.5 U of Proteinase K (right) at 37°C for 1 h, followed by SDS-PAGE and staining with Coomassie Blue. Proteinase K treatment resulted in loss of all visible protein bands.

**Supplementary Figure 3.**
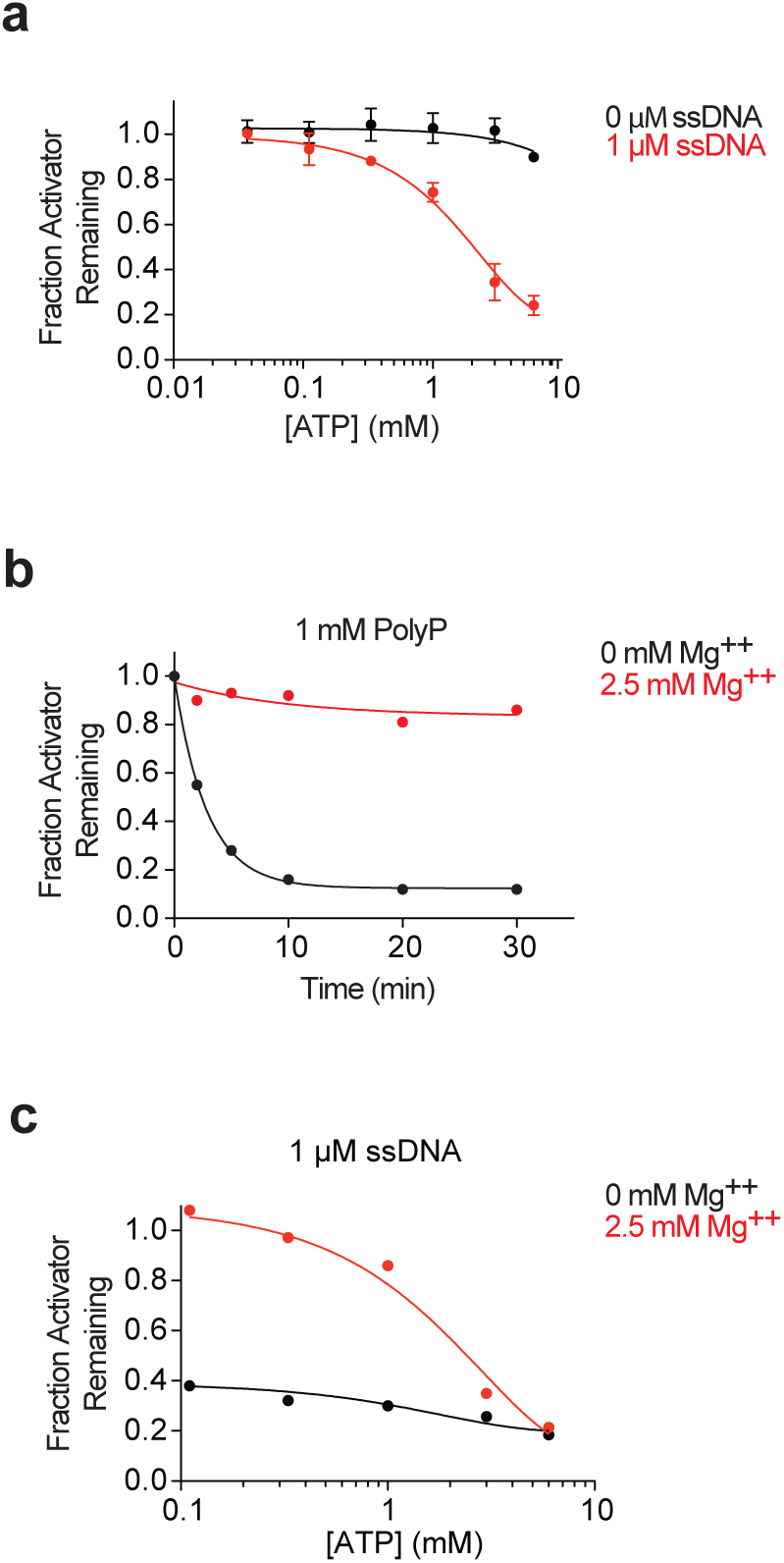
ATP stimulates nucleic-acid-dependent dissociation by sequestering magnesium ions. **a.** ATP alone is not sufficient for dissociation. Varying ATP concentrations were tested in Cdh1 dissociation assays. Reactions were performed with buffer alone (black) or 1 μM 75mer ssDNA (red) in the presence of 2.5 mM MgCl_2_. **b.** Magnesium inhibits the effects of polyphosphate. Cdh1 dissociation reactions were performed using polyphosphate (1 mM total PO_4_^-^) in the absence (black) or presence (red) of 2.5 mM MgCl_2_. **c.** ATP stimulates activator dissociation only when magnesium is present. Cdh1 dissociation reactions were performed using 1 μM 75mer ssDNA. ATP was serially diluted and added to reactions in the absence (black) or presence (red) of 2.5 mM MgCl_2_.

**Supplementary Figure 4.**
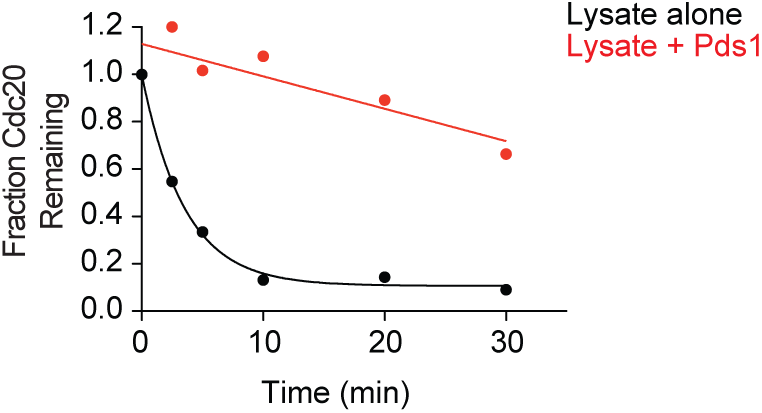
Pds1 binding inhibits Cdc20 dissociation. Cdc20 dissociation reactions were performed in the presence of yeast lysate (2.5 mg/ml) and 5 mM ATP, in the absence (black) or presence (red) of 10 μM Pds1 fragment (aa 1-110). His-tagged Pds1 fragment was expressed in bacteria and purified using a nickel column.

**Supplementary Table 1.**
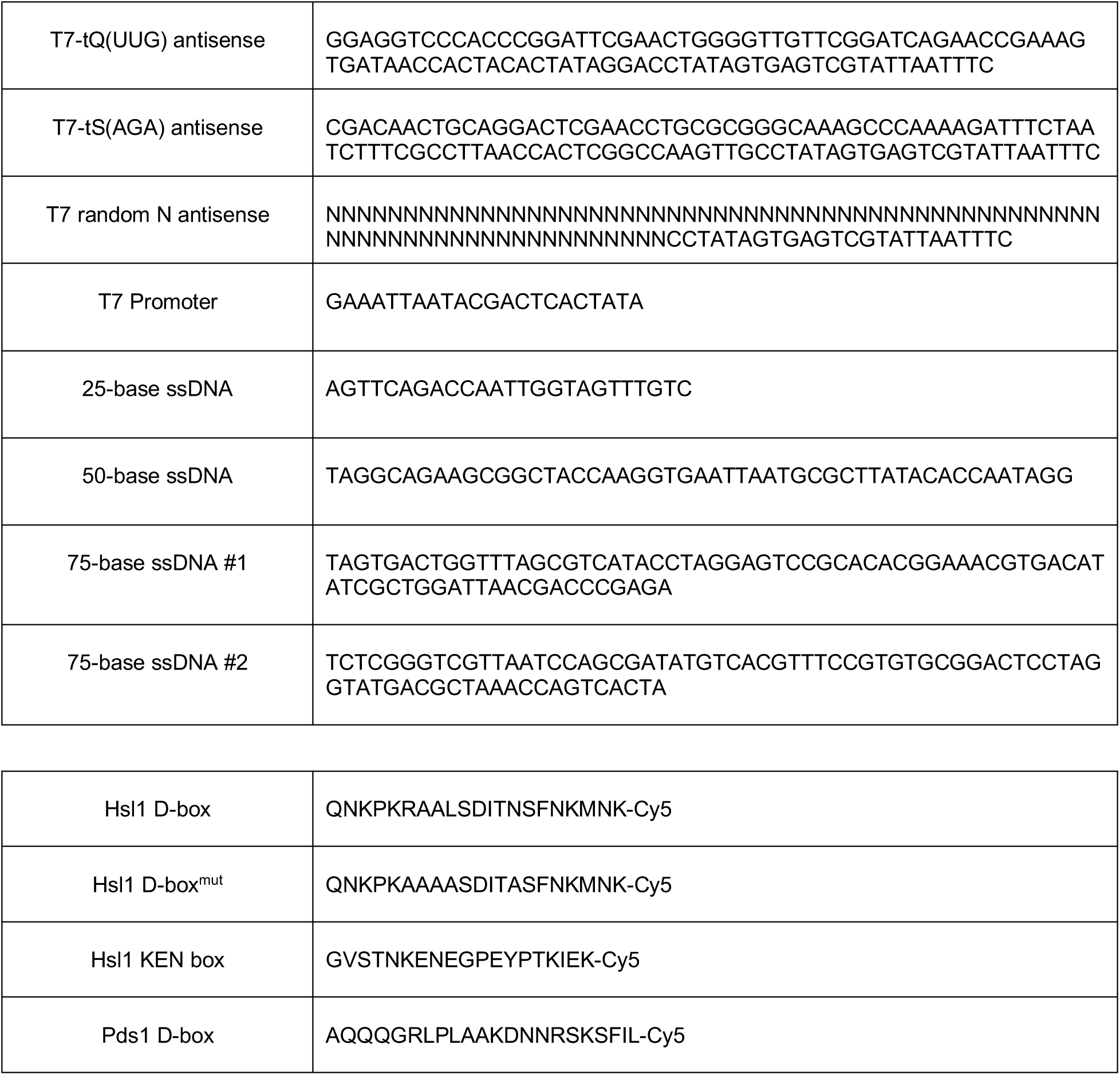
Nucleic acid and peptide sequences used in this study. T7 antisense sequences were annealed with T7 promoter sequence and used for transcription of the indicated RNAs *in vitro*.

